# Construction, validation and application of nocturnal pollen transport networks in an agro-ecosystem: a comparison using microscopy and DNA metabarcoding

**DOI:** 10.1101/325084

**Authors:** Callum J. Macgregor, James J.N. Kitson, Richard Fox, Christoph Hahn, David H. Lunt, Michael J.O. Pocock, Darren M. Evans

## Abstract

1. Moths are globally relevant as pollinators but nocturnal pollination remains poorly understood. Plant-pollinator interaction networks are traditionally constructed using either flower-visitor observations or pollen-transport detection using microscopy. Recent studies have shown the potential of DNA metabarcoding for detecting and identifying pollen-transport interactions. However, no study has directly compared the realised observations of pollen-transport networks between DNA metabarcoding and conventional light microscopy.
2. Using matched samples of nocturnal moths, we construct pollen-transport networks using two methods: light microscopy and DNA metabarcoding. Focussing on the feeding mouthparts of moths, we develop and provide reproducible methods for merging DNA metabarcoding and ecological network analysis to better understand species-interactions.
3. DNA metabarcoding detected pollen on more individual moths, and detected multiple pollen types on more individuals than microscopy, but the average number of pollen types per individual was unchanged. However, after aggregating individuals of each species, metabarcoding detected more interactions per moth species. Pollen-transport network metrics differed between methods, because of variation in the ability of each to detect multiple pollen types per moth and to separate morphologically-similar or related pollen. We detected unexpected but plausible moth-plant interactions with metabarcoding, revealing new detail about nocturnal pollination systems.
4. The nocturnal pollination networks observed using metabarcoding and microscopy were similar, yet distinct, with implications for network ecologists. Comparisons between networks constructed using metabarcoding and traditional methods should therefore be treated with caution. Nevertheless, the potential applications of metabarcoding for studying plant-pollinator interaction networks are encouraging, especially when investigating understudied pollinators such as moths.

## Introduction

Species interaction networks, which describe the presence and strength of interspecific interactions within ecosystems (Montoya *et al.*, 2006), are an important tool in understanding and conserving ecosystem processes and functioning (Tylianakis *et al.*, 2010). Currently, there is considerable interest in pollination networks, due to ongoing global declines in pollinating insects (Potts *et al.*, 2010) and their role in reproduction of both wild plants and crops (Klein *et al.*, 2007; Ollerton *et al.*, 2011).

Many flower-visiting animals are not effective pollinators, and proving the existence of an effective pollination interaction is labour-intensive (King *et al.*, 2013). Consequently, proxies for pollination are often used to construct plant-pollinator interaction networks, which cannot strictly be referred to as pollination networks. A commonly-used proxy is flower-visitation, recorded by directly observing animals visiting flowers. This is effective for daytime sampling, but is challenging to apply to nocturnal pollinators, such as moths (Lepidoptera; Macgregor *et al.*, 2015), because observations are difficult and may be biased if assisted by artificial light. This may explain why plant-pollinator network studies frequently omit nocturnal moths, even though moths are globally relevant pollinators (Macgregor *et al.*, 2015).

An alternative to direct observation is detecting pollen transport, by sampling and identifying pollen on the bodies of flower-visiting animals; this approach has been used in several previous studies of nocturnal pollination by moths (Devoto *et al.*, 2011; Banza *et al.*, 2015; Knop *et al.*, 2017; Macgregor *et al.*, 2017a). By analysing pollen transport, flower-visits where no pollen is received from the anthers are excluded (Pornon *et al.*, 2016). This approach can detect more plant-pollinator interactions with lower sampling effort than flower-visitor observations (Bosch *et al.*, 2009). Studies of pollen transport also permit unbiased community-level sampling of interactions without requiring decisions about distribution of sampling effort among flower species, as each pollinator carries a record of its flower-visiting activities in the pollen on its body (Bosch *et al.*, 2009). Traditionally, pollen identification is undertaken using light microscopy with a reference collection of known species (e.g. Devoto *et al.*, 2011). However, identifications made by microscopy can be ambiguous, especially when distinguishing related species (Galimberti *et al.*, 2014). Accurate, reproducible identification of pollen sampled from pollinators is necessary to ensure plant-pollinator networks are free from observer bias.

A recent alternative to microscopy is DNA metabarcoding: high-throughput sequencing of standard reference loci from communities of pooled individuals (Cristescu, 2014). It offers possibilities to detect interspecific interactions, including plant-pollinator interactions (Evans *et al.*, 2016), and methods are rapidly improving, permitting greater accuracy in species identification (Bell *et al.*, 2016a) for reducing costs (Kamenova *et al.*, 2017). Studies using metabarcoding have identified pollen sampled from honey (Hawkins *et al.*, 2015; de Vere *et al.*, 2017) and directly from bees (Galimberti *et al.*, 2014) and flies (Galliot *et al.*, 2017), and constructed plant-pollinator networks (Bell *et al.*, 2017; Pornon *et al.*, 2017). DNA sequences have confirmed identities of single pollen grains sampled from moths (Chang *et al.*, 2018), but no study has applied metabarcoding to nocturnal pollen-transport by moths, where pollen-transport approaches may be most valuable, given the paucity of existing knowledge about moth-plant pollination interactions. Metabarcoding reveals more plant-pollinator interactions than direct flower-visitor observations (Pornon *et al.*, 2016, 2017), but it is unclear whether this is purely because pollen-transport approaches detect interactions more efficiently than flower-visitation approaches (Bosch *et al.*, 2009) or whether metabarcoding offers specific additional benefits. Use of a metabarcoding approach is often justified by the labour-intensive nature of microscopy-based approaches and the level of expertise required to identify pollen morphologically (e.g. de Vere *et al.*, 2017). It is frequently suggested that metabarcoding increases the level of species discrimination compared to traditional approaches (Bell *et al.*, 2017). Crucially, despite this assertion, no study has directly compared metabarcoding to traditional microscopy for assessing pollen transport. It is therefore unknown whether, in studies using a pollen-transport approach, the choice of detection method (light microscopy or DNA metabarcoding) can alter the realised observations of plant-pollinator interactions.

In this study, we used matched samples of moths to construct nocturnal pollination networks using two methods: DNA metabarcoding, and the traditional light microscopy approach; and compared the observed networks, considering the quantity and nature of the interactions detected and the properties of the networks themselves. We sampled moths in a UK agro ecosystem, as our previous study suggests that moths may have greater importance as pollinators in such systems than generally thought (Macgregor *et al.*, 2017a). Accordingly, we developed existing pollen-metabarcoding protocols to enable detection of pollen transported by moths, and integrated molecular advances with ecological network analysis to provide a reproducible methodology for the improved study of species-interactions. By providing detailed descriptions of our methods (dx.doi.org/10.17504/protocols.io.mygc7tw, Appendix S1) and archiving all bioinformatic and statistical code (dx.doi.org/10.5281/zenodo.1169319), we present a framework for future studies of pollination networks using metabarcoding. We discuss the advantages and disadvantages of each method for assessment of pollen transport by moths and other pollinator taxa, current limitations and future research directions.

## Materials and methods

### Field sampling

We sampled moths, using light-traps, from four locations in a single farmland site in the East Riding of Yorkshire, UK (53°51’44" N 0°25’14" W), over eight nights between 30th June and 19th September 2015 (Table S1; full details in Appendix S1). Moths were euthanised and retained individually. As both pollen-sampling methods are destructive, it was impossible to directly compare sensitivity by sampling pollen from the same individual moth with both methods. Instead, we created two matched sub-samples of moths, each containing the same set of species, and the same number of individuals of each. Pollen-transport by each sub-sample was analysed using one method (Fig. 1). With both methods, we restricted pollen sampling to the proboscis, because most moth species coil their proboscides unless actively feeding (Krenn, 1990). Therefore, the proboscis is unlikely to experience cross contamination of pollen through contact with other moths (e.g. whilst in the moth-trap), and pollen held on the proboscis is probably the result of a flower-visitation interaction.

**Figure 1:**
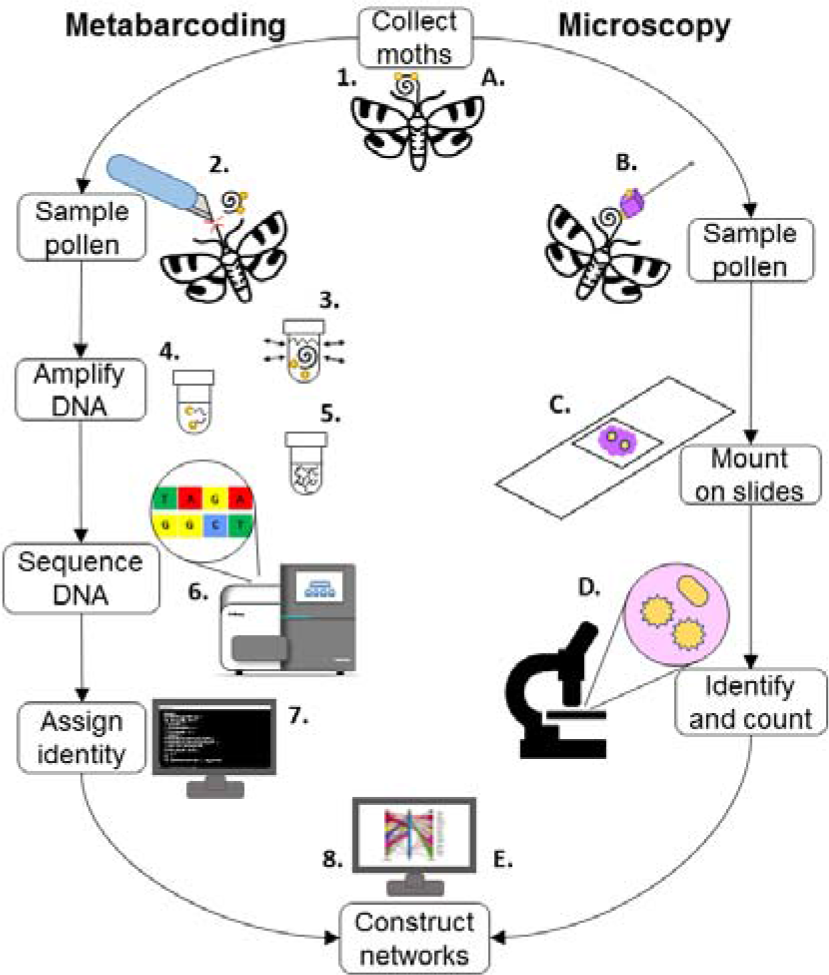
visual summary of the two methods applied to detect and identify pollen transport by moths. Full methods are in Appendix S1. For metabarcoding, the steps shown are: 1. Field sampling of moths. 2. Excise proboscis. 3. Remove pollen by shaking. 4. Extract DNA by HotSHOT method. 5. Amplify DNA by 3-step PCR protocol. 6. Sequence DNA. 7 Assign DNA sequence identities. 8. Analyse interactions and construct networks. For microscopy, the steps shown are: A. Field sampling of moths. B. Swab proboscis with fuchsin-stained gel. C. Mount gel on microscope slide. D. Identify and count pollen under microscope. E. Analyse interactions and construct networks.

### Method 1: light microscopy

A standard approach for pollen sampling was applied (Beattie, 1972), in which 1 mm^3^ cubes of fuchsin jelly were used to swab pollen from the proboscides of moths, and the pollen examined under a light microscope at 400x magnification. Pollen morphotypes were identified using a combination of keys (Moore *et al.*, 1994; Kapp *et al.*, 2000) and knowledge of likely insect-pollinated plant taxa. Morphotypes (equivalent to operational taxonomic units, OTUs) represented groupings that could not be unambiguously separated to a lower taxonomic level, and might have contained pollen from multiple species.

### Method 2: DNA metabarcoding

Protocols for DNA extraction, amplification and sequencing are fully described in Appendix S1 and archived online (dx.doi.org/10.17504/protocols.io.mygc7tw). In brief, the protocols were as follows. Moth proboscides were excised using a sterile scalpel. Pollen was removed from each proboscis by shaking for 10 minutes in HotSHOT lysis reagent (Truett *et al.*, 2000) at 2000 rpm on a Variomag Teleshake plate shaker (Thermo Scientific, Waltham, MA). The proboscis was removed using sterile forceps, and the DNA extraction procedure completed on the remaining solution following Truett *et al.* (2000). Extracted DNA was amplified using a three-step PCR nested tagging protocol (modifed from Kitson *et al.*, n.d. in press; see Appendix S1). We amplified a custom fragment of the *rbcL* region of chloroplast DNA, which has been previously used for metabarcoding pollen (Hawkins *et al.*, 2015; Bell *et al.*, 2017) and has a comprehensive reference library for the Welsh flora, representing 76% of the UK flora (de Vere *et al.*, 2012), available on the International Nucleotide Sequence Database Collaboration (http://www.insdc.org/; GenBank). We used two known binding sites for reverse primers, rbcL-19bR (Hofreiter *et al.*, 2000) and rbcLr506 (de Vere *et al.*, 2012), to produce a working forward and reverse universal primer pair, rbcL-3C (rbcL-3CF: 5‘-CTGGAGTTCCGCCTGAAGAAG-3’; rbcL-3CR: 5‘-AGGGGACGACCATACTTGTTCA-3’). Primers were validated by successful amplification of DNA extracts from 23/25 plant species (Table S2). Sequence length varied widely (median: 326 base pairs (bp), range: 96-389 bp); fragments shorter than 256 bp generally had no match on GenBank. Six control samples were used to monitor cross-contamination between wells (Table S3).

Amplified DNA was sequenced on an Illumina MiSeq, using V2 chemistry. Taxonomic assignment of MiSeq output was conducted using the metaBEAT pipeline, version 0.97.7 (https://github.com/HullUni-bioinformatics/metaBEAT). For reproducibility, all steps were conducted in Jupyter notebooks; all bioinformatic and statistical code used in this study is archived online (dx.doi.org/10.5281/zenodo.1169319) and procedures are explained in full in Appendix S1. Taxonomic assignment of sequences was conducted within metaBEAT based on a BLAST Lowest Common Ancestor approach implemented in MEGAN (Huson *et al.*, 2007). We chose to conduct taxonomic assignment with BLAST because it is among the most widely-used taxonomic assignment tools, and blastn specifically has a proven capacity to discriminate between UK plant species using the *rbcL* locus (de Vere *et al.*, 2012). We used a curated database of reference sequences from plausibly-present plant species previously recorded in the vice-county of South-east Yorkshire (reference list of species archived at dx.doi.org/10.5281/zenodo.1169319).

To eliminate the risk of cross-well contamination, we established a threshold for minimum read depth of 50 reads, per assignment, per well. The maximum read depth in any negative control well was 47, and the maximum read depth in any positive control well of sample assignments was 33 (Table S3). Therefore, this threshold was adequate to remove sample reads from positive and negative controls. Within each well, any assignment with a read depth below 50 was reset to 0 prior to statistical analysis; this resulted in some plant OTUs being removed entirely from the dataset (however, these OTUs are indicated in Table 1).

**Table 1:**
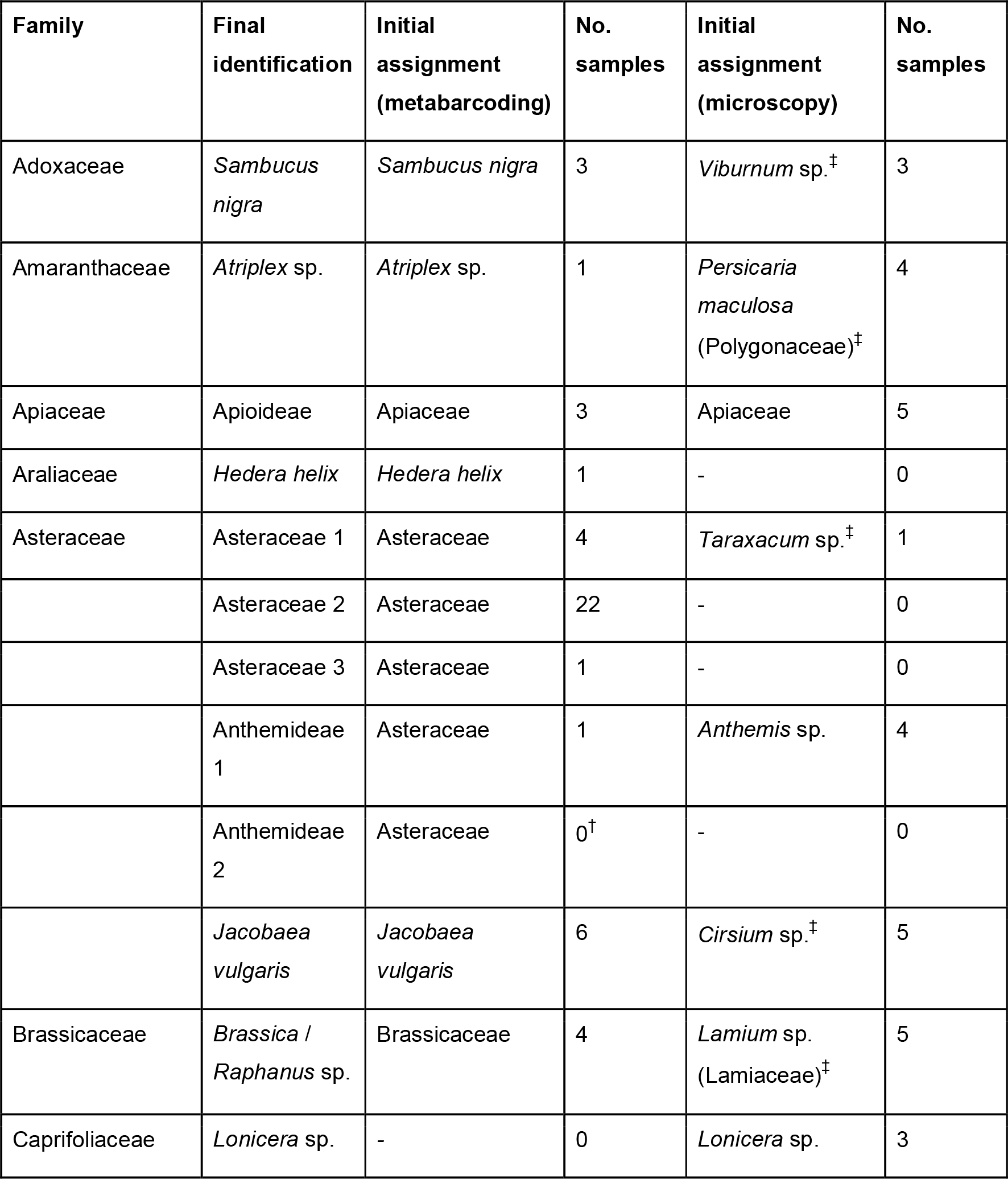
harmonised plant OTUs identified by metabarcoding and microscopy. In column 4, ^†^ indicates an assignment initially identified by metabarcoding, but failing to meet the minimum read depth threshold in any sample (Table S7). In column 5, ^‡^ indicates an assignment that was re-identified by comparison to pollen of species identified by metabarcoding.

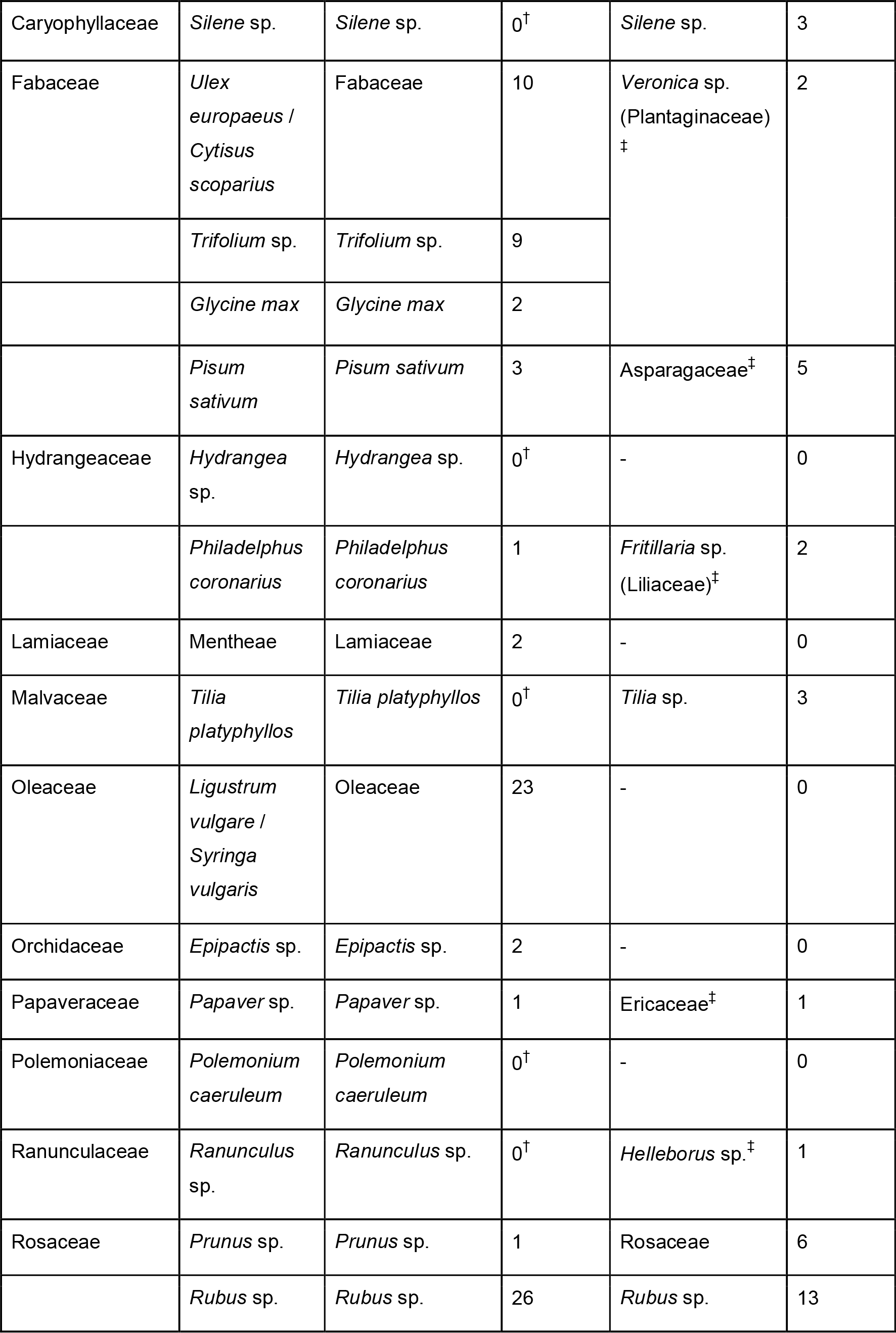

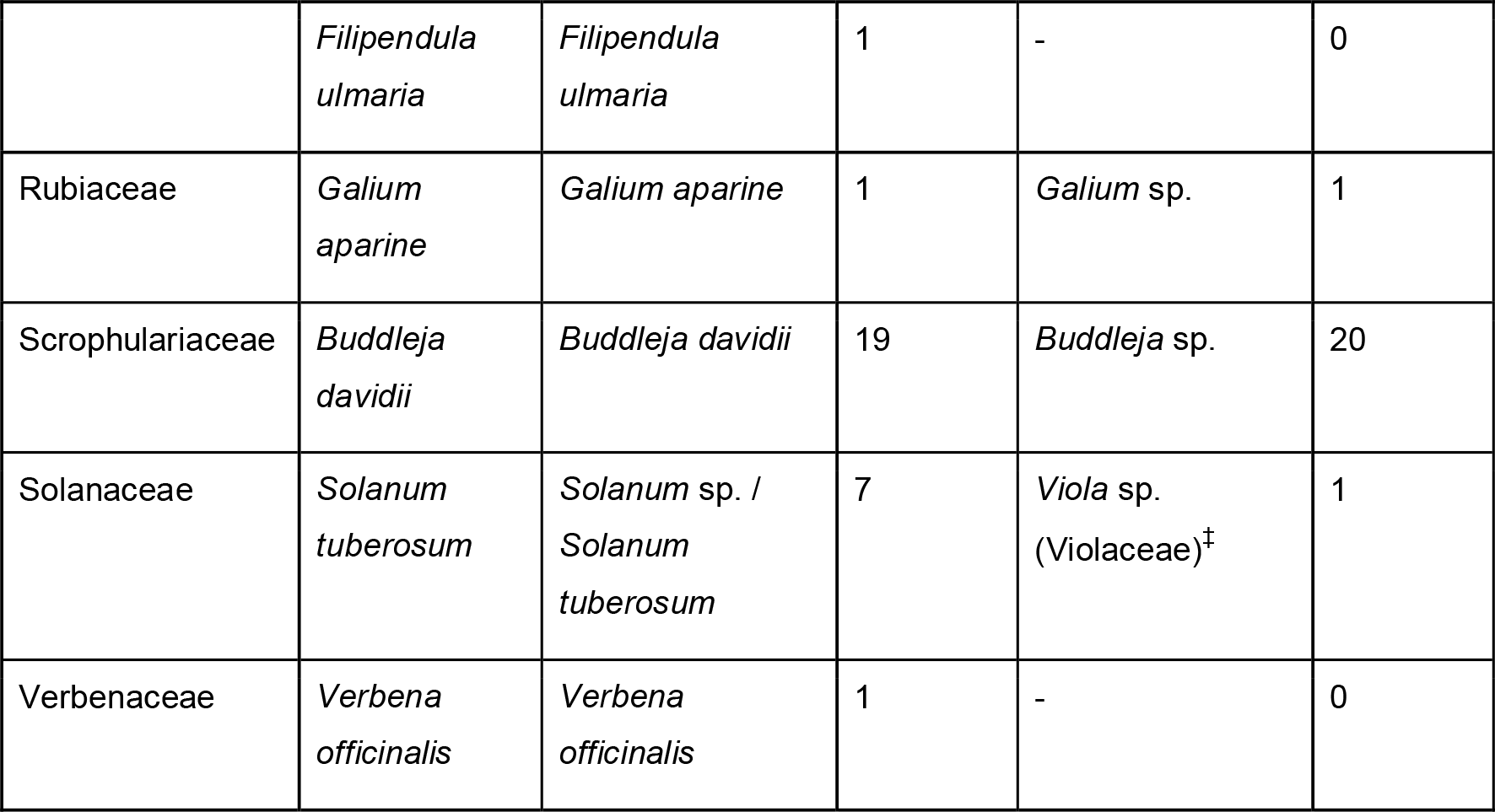

**Table 2:** Summary of basic interaction data for each method. The samples were duplicate subsets of the total sample, and each comprised 311 individuals of 41 species. Plant types for metabarcoding were operational taxonomic units (OTUs; identified by a BLAST search against a curated reference database) and for microscopy were morphotypes (identified using identification keys). Percentages in brackets are of the relevant sub-sample.

|  | Metabarcoding | Microscopy |
| --- | --- | --- |
| No. pollen-carrying moths | 107 (34.4%) | 70 (22.5%) |
| No. pollen-carrying species | 15 (36.6%) | 17 (41.5%) |
| No. plant types identified | 26 | 20 |
| Plant types initially identified to species level | 11 (42.3%) | 1 (5%) |
| Plant types initially identified to at least genus level | 17 (65.4%) | 16 (80%) |
| Plant types detected on one moth only | 10 (38.5%) | 5 (25%) |
| No. moths carrying pollen from >1 plant types | 36 (11.6%) | 13 (4.2%) |
| No. unique interactions (total no. interactions) | 62 (155) | 52 (88) |

### Curation of data

We harmonised the plant identifications from each method (OTUs from metabarcoding and morphotypes from microscopy) to produce a single list of plants consistent across both methods (Table 1). Specifically, for metabarcoding, we revised family-level assignments made by BLAST, inspecting the range of species-level matches to identify clear taxonomic clusters within the families. For microscopy, we attempted to re-identify pollen morphotypes using images of pollen from species identified by metabarcoding for additional reference (see Appendix S1). Microscopic photographs of pollen were sourced from two online repositories of pollen images: Pollen-Wiki (http://pollen.tstebler.ch/MediaWiki/index.php?title=Pollenatlas) and the Pollen Image Library (http://www-saps.plantsci.cam.ac.uk/pollen/index.htm).

### Comparison of methods and statistical analysis

We tested for differences between the two identification methods, examining whether sampling method affected the likelihood of detecting (i) pollen on individual moths; (ii) more than one pollen species on individuals; (iii) pollen on moth species (individuals combined); and whether sampling method affected the number of pollen types detected (iv) per individual moth; and per moth species, using (v) observed richness and (vi) true richness estimated using the Chao2 estimator (Chao, 1987). We used generalised linear mixed-effects models (GLMMs), with sampling method as a fixed effect. In individual-level analyses, we used date/light-trap combination (‘trap ID’) as a random effect, whilst in species-level analyses, we used moth species as a random effect to treat the data as pairs of observations (one observation, per method, per moth species). We tested significance of fixed effects using either Likelihood Ratio Tests or Type III ANOVA, depending on error distribution. Analysis was carried out with R version 3.3.2 (R Core Team, 2016); all code is archived at dx.doi.org/10.5281/zenodo.1169319.

### Sampling completeness and networks

For both methods, we estimated sampling completeness of interactions, following Macgregor *et al.* (2017b). For each method, we estimated the total number of pollen types (interaction richness) for each insect species with the Chao2 estimator (Chao, 1987), using the R package vegan (Oksanen *et al.*, 2015). We calculated interaction sampling completeness for each species as 100*(observed interactions)/(estimated interactions) for each species. Finally, we calculated the mean interaction sampling completeness of all species, weighted by estimated interaction richness of each species.

We constructed pollen-transport networks from the interaction data. We used presence of interactions between individual moths and plant taxa, rather than strength of individual interactions, because read depth (metabarcoding) and pollen count (microscopy) are not equivalent. We measured interaction frequency by counting interactions across all individuals in each moth species; interaction frequency correlates positively with true interaction strength in mutualistic networks (Vázquez *et al.*, 2005). We calculated several quantitative metrics, as follows, to describe the diversity and specialisation of interactions forming each network. Improved detection of interactions could increase the complexity of the network, so we calculated two measures of network complexity: linkage density (average no. links per species) and connectance (proportion of possible interactions in the network that are realized). Likewise, improved detection of plant species with the same set of pollinator species could alter consumer-resource asymmetry and perceived specialization of species in the network, so we calculated H2’ (a frequency-based index that increases with greater specialization), generality of pollinators, and of plants (average no. links to plant species per pollinator species, and *vice versa*). Finally, the resilience of the network to cascading species loss may be influenced by its complexity (Dunne *et al.*, 2002), so we measured the robustness of each network (mean robustness across 1000 bootstrapped simulations of pollinator species loss). For comparison, we repeated all network analyses with plant identities aggregated at family-level, because the methods might differ in their ability to distinguish closely-related species. Networks were analysed using the package bipartite (Dormann *et al.*, 2009) and plotted using Food Web Designer 3.0 (Sint & Traugott, 2016). As we could only construct one network for each method, we recorded obvious differences between the metrics for each network but could not statistically assess the significance of those differences.

## Results

### Summary

In total, we caught 683 moths of 81 species, generating two matched sub-samples, each containing 311 moths of 41 species (Table S4). We detected pollen on 107 individual moths with metabarcoding (34% of the sub-sample) and 70 (23%) with microscopy. We initially identified 20 plant morphotypes in the microscopy sample and 25 OTUs in the metabarcoding sample (Table 1). After harmonising these we recorded 33 plant identities (at varying taxonomic resolution), of which 18 were detected with both methods, 11 with metabarcoding only (including three which failed to meet the minimum read depth threshold in any sample), and four by microscopy only.

### Statistical comparisons between methods

Metabarcoding was significantly more likely than microscopy to detect pollen (Fig. 2) on individual moths (◻^2^ = 10.95, *P* < 0.001), and to detect more than one pollen type on individual moths (◻^2^ = 12.00, *P* < 0.001). However, with non-pollen-carrying moths excluded, the methods did not differ in the number of pollen types detected per individual moth (◻^2^ = 1.12, *P* = 0.290). With data aggregated per moth species, the methods did not differ in the likelihood of detecting pollen (◻^2^ = 0.37, *P* = 0.545), but metabarcoding detected significantly more pollen types per moth species (◻^2^ = 18.09, *P* < 0.001); this difference was non significant when the estimate of true interaction richness was used (◻^2^ = 3.62, *P* = 0.057; Table S5).

**Figure 2:**
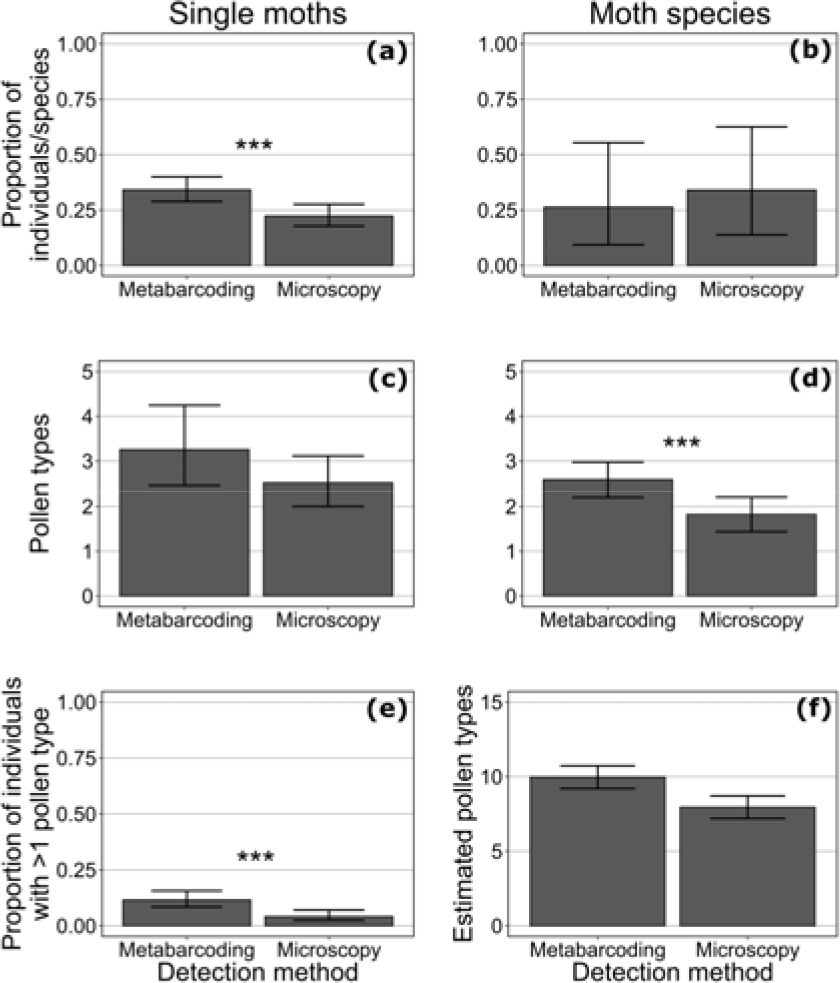
**comparisons between DNA metabarcoding and microscopy approaches** of: proportion of (a) individual moths and (b) moth species found to be carrying pollen; number of pollen types detected for (c) individual moths and (d) moth species; proportion of individual moths carrying more than one pollen type (e); and estimated number of pollen types per moth species (f). For (c), (d) and (f). only pollen-carrying individuals and moth species were included. Significance indicates Likelihood Ratio Test for detection method in GLMMs (* : *P* <0.05; ** : *P* <0.01; *** *P* <0.001). Error bars show 95% confidence intervals.

### Construction and analysis of networks

For each method, we constructed a quantitative pollen-transport network (Fig. 3). The estimated sampling completeness of interactions was higher for the microscopy network (75.7%) than the metabarcoding network (43.2%). Some network metrics differed markedly between the two methods (Fig. 4), though no statistical comparison was appropriate. Specifically, linkage density and generality of pollinators were higher in the metabarcoding network than the microscopy network, but all other metrics were similar. With plant assignments aggregated at family level, the metabarcoding network had higher generality of pollinators and lower generality of plants than the microscopy network (Table S6).

**Figure 3:**
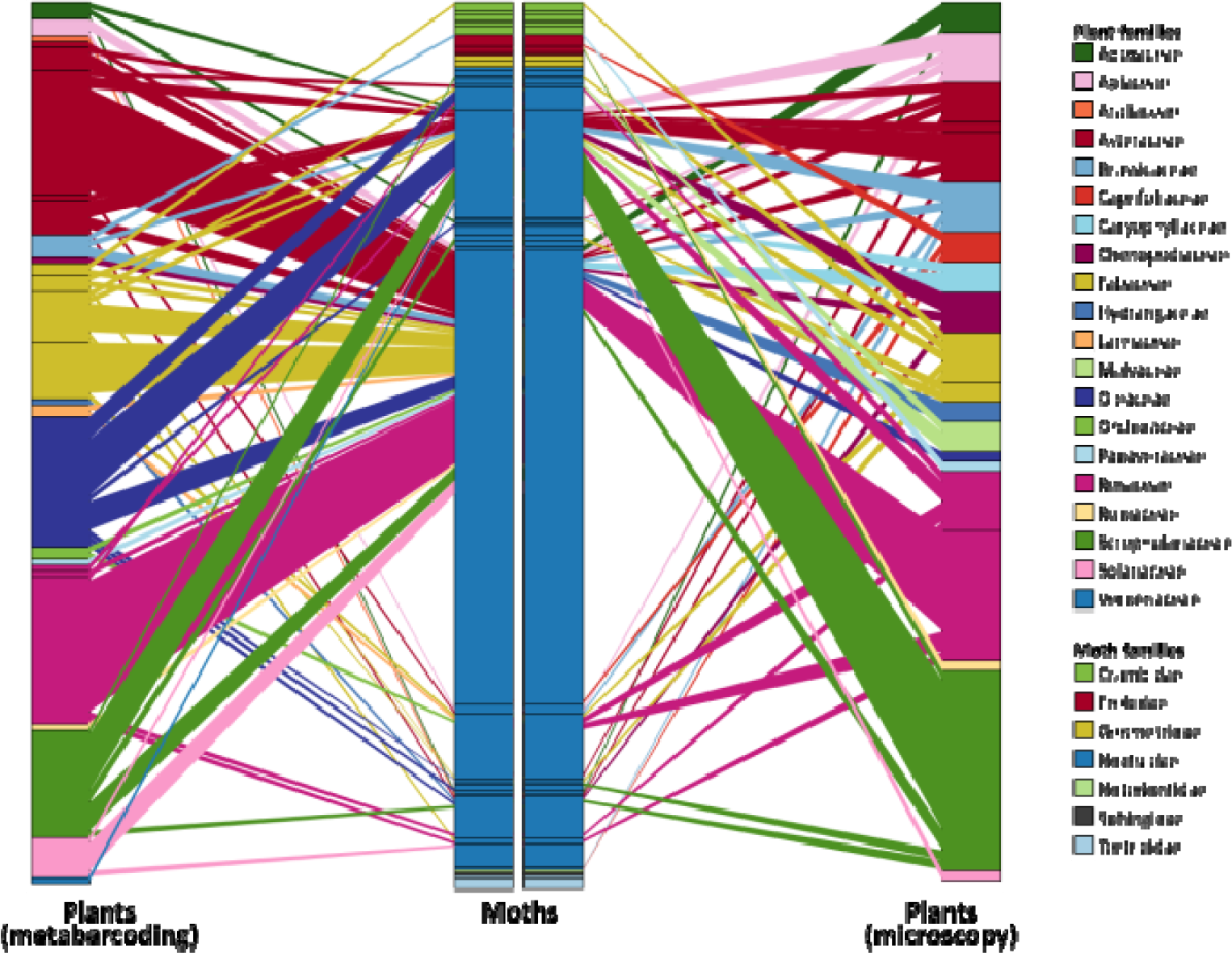
networks constructed using DNA metabarcoding and microscopy from replicated, matched samples of moths. Species are colour-coded by family (see key); families appear from top to bottom in the order listed. For moths, bar height indicates relative species abundance, and link width indicates number of individuals carrying pollen of each plant species. For plants, bar height indicates number of individual moths on which each pollen type was detected, and link width indicates proportion of those moths belonging to each moth species.

## Discussion

### Methodological comparison

Our realised observations of the plant-pollinator system were generally similar between the DNA-based (metabarcoding) and microscopy-based methods for detecting and identifying pollen-transport by moths, but nonetheless showed some key differences. Metabarcoding detected more pollen OTUs in total than microscopy, detected pollen on a greater proportion of individual moths, and was more likely to detect multiple pollen OTUs on a moth. When moths were aggregated to species level, metabarcoding detected more pollen types in total per moth species.

We observed differences between the networks detected by each method, which can be attributed to metabarcoding detecting more separate species within some plant families, and detecting interactions with more plant families per pollinator species. This is revealed by the higher generality of pollinators in the fully-resolved metabarcoding network than its equivalent microscopy network, and the lesser increase in generality of pollinators, combined with lower generality of plants, in the family-level metabarcoding network than its equivalent (Fig. 4). Additionally, linkage density was higher for metabarcoding than microscopy in the fully-resolved networks, but not in the family-level networks (Fig. 4).

**Figure 4:**
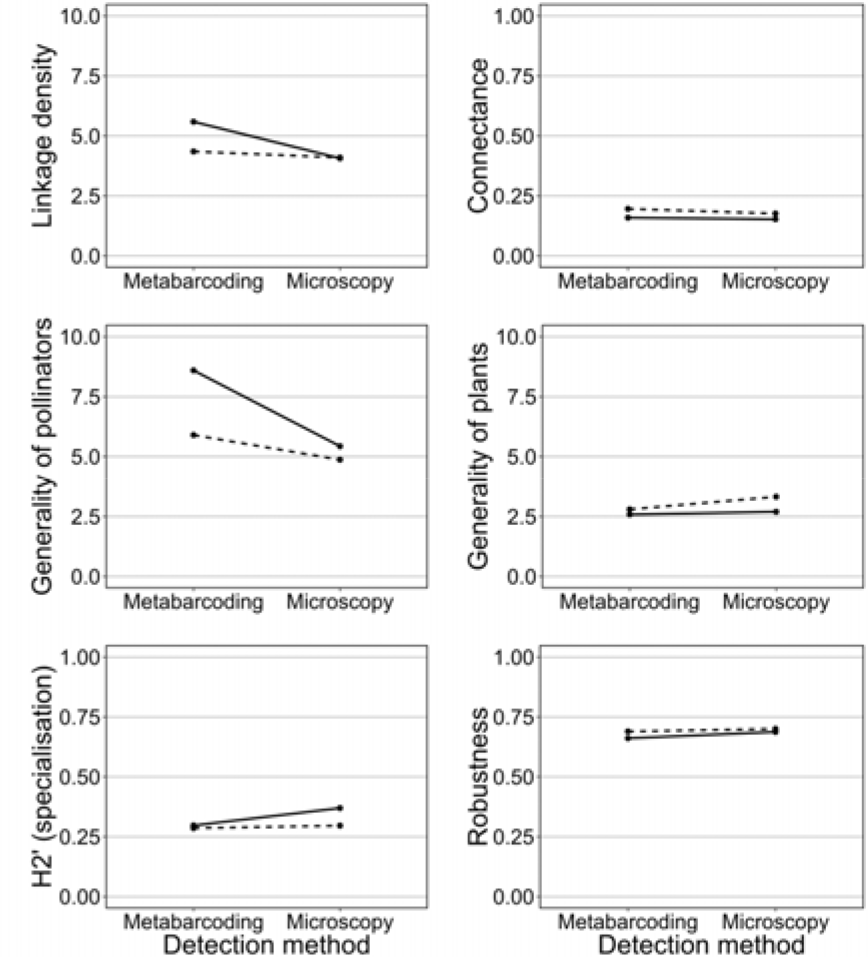
network metrics calculated for each detection method. (Table S6). Solid lines connect metrics for fully-resolved data, dashed lines connect metrics when plant species were aggregated at the family level.

Estimated sampling completeness of interactions differed conspicuously between networks (Table S6). Despite containing more interactions, the metabarcoding network was estimated to be less completely sampled than the microscopy network. This is probably because metabarcoding detected more ‘rare’ interactions (‘singletons’, detected only once), being more effective at distinguishing morphologically-similar pollen. This would result in a higher ratio of singletons to doubletons (interactions detected twice) and therefore a proportionally greater estimated value of interaction richness. This demonstrates that sampling method can substantially affect estimation of sampling completeness of interactions in network studies.

### Pollen transported by moths

We identified several plants using metabarcoding that were not initially identified as the same species by microscopy. Because many plants have morphologically-similar pollen, we conservatively chose not to identify novel moth-flower associations by microscopy unless the identification was unambiguous. Among the plants initially identified only by metabarcoding were species for which moths were not previously recorded in the literature as pollinators or flower-visitors (Macgregor *et al.*, 2015), highlighting that much is still unknown about pollination by moths. Some of these fitted the moth-pollination ‘syndrome’ (Grant, 1983), being white and fragrant: *Sambucus nigra* (Adoxaceae), *Philadelphus coronarius* (Hydrangeaceae)*, Filipendula ulmaria* (Rosaceae) and *Ligustrum vulgare* (Oleaceae; though not *Syringa vulgaris*, not separable in this study). However, others did not and are typically associated with other pollinators: for example, *Polemonium caerulum* (Polemoniaceae) and *Trifolium* spp. (Fabaceae) are visited by bees (Palmer-Jones *et al.*, 1966; Zych *et al.*, 2013), *Verbena officinalis* (Verbenaceae) is most likely visited by bees and butterflies (Perkins *et al.*, 1975), whilst species of *Epipactis* (Orchidaceae) are generalist, with previously-known visitors including diurnal Lepidoptera (Jakubska-Busse & Kadej, 2011).

We found pollen from plants that, in this region, are chiefly associated with domestic gardens, including two species of Hydrangeaceae, species from the tribe Mentheae (Lamiaceae; includes many species grown as culinary herbs, though wild species might also have occurred), *Buddleja davidii* (Scrophulariaceae; though a railway ran adjacent to the farm and *B. davidii* is widely naturalised along railways in the UK) and *Verbena officinalis* (Verbenaceae). Individual moths may have carried pollen several hundred metres from the closest gardens to the field site. This provides new evidence to support previous suggestions that moths could play an important role in providing gene flow among plant populations at the landscape-scale (Miyake & Yahara, 1998; Young, 2002; Barthelmess *et al.*, 2006), and even at continental scales for species of moths that undergo long-distance migrations (Chang *et al.*, 2018). Such gene flow could provide benefits from nocturnal pollination even to plant species that are primarily diurnally-pollinated and not pollination-limited.

Finally, we detected several insect-pollinated crop species (only some of which require pollination for crop production): specifically, soybean *Glycine max* and pea *Pisum sativum* (Fabaceae), potato *Solanum tuberosum* (Solanaceae), and *Brassica/Raphanus* sp. (includes oil-seed rape; Brassicaceae). Floral phenology suggests *Prunus* sp. (Rosaceae) was likely to be cherry *(P. avium, P. cerasus* or a hybrid) rather than wild *P. spinosa.* Similarly, *Rubus* sp. (Rosaceae) could have been wild blackberry (matching to *R. caesius, R. plicatus* and *R. ulmifolius)* but also matched raspberry *R. idaeus.* There is currently an extreme paucity of evidence in the existing global literature to support a role of moths in providing pollination services by fertilizing economically-valuable crops (Klein *et al.*, 2007; Macgregor *et al.*, 2015). Although our findings do not prove that any of the crops recorded receive significant levels of nocturnal pollination by moths, they do highlight a vital and urgent need for further research into the potential role of moths as pollinators of agricultural crop species.

### Current methodological limitations

We identified limitations with both methods, relating to the accuracy and taxonomic resolution of pollen identification and the non-quantitative interaction data they generated.

Firstly, there was little initial overlap between identifications made by each method (of 20 initial assignments from microscopy and 25 from metabarcoding, only 3 plant identifications were shared between methods at genus- or species-level). Because we applied the methods to separate samples of moths, some differences were expected between the pollen species transported. In two cases (*Silene* and *Tilia*), species identified by microscopy were discarded from the metabarcoding assignments by application of the 50-reads threshold. Both species had very low abundance in microscopy samples (<20 pollen grains per sample), suggesting precautions against cross-sample contamination with metabarcoding might mask detection of low-abundance pollen. The remaining mismatches were most probably misidentifications by one or other method. Using images of pollen from species identified by metabarcoding as a reference for microscopy, we re-identified several pollen morphotypes, increasing agreement between the methods (19 identifications matched across methods, of which 10 were at genus- or species-level; Table 1). Misidentifications were arguably more likely under microscopy than metabarcoding, due to the conservative approach used when applying BLAST and the difficulty of unambiguously identifying pollen by microscopy.

Secondly, several assignments made with metabarcoding were not resolved beyond family-level. Although *rbcL* is a popular marker region for plant barcoding (Hawkins *et al.*, 2015) and has been shown to identify over 90% of Welsh plants to at least genus-level using blastn (de Vere *et al.*, 2012), interspecific sequence diversity within *rbcL* is nonetheless extremely low within some families (e.g. Apiaceae; Liu *et al.*, 2014). In some cases, reference sequences from multiple genera did not differ across our entire fragment, leading BLAST to match query sequences to species from several genera with equal confidence. Such instances could not have been further resolved using our fragment, even by alternative assignment methods. Sequencing a longer fragment might increase interspecific sequence variation; improvements in sequencing technology may facilitate accurate sequencing of such longer amplicons (Hebert *et al*., 2017). Using another locus than *rbcL* might improve taxonomic resolution; loci including ITS2 and *matK* are also used to metabarcode pollen (Bell *et al.*, 2016b). Sequencing two or more of these loci simultaneously might also improve assignment resolution (de Vere *et al.*, 2012), though at greater cost.

Thirdly, some studies have weighted interactions in networks using the number of pollen grains transported, as a proxy for interaction strength (e.g. Banza *et al.*, 2015). This approach is impossible with metabarcoding, as the number of pollen grains in a sample does not correlate with read depth (Pornon *et al.*, 2016), and metabarcoding cannot definitively distinguish pollen from other sources of plant DNA (e.g. residual nectar on mouthparts). However, an insect’s pollen load also may not be a true indicator of its efficacy as a pollinator (Ballantyne *et al.*, 2015); pollinator effectiveness differs between pairwise interactions through variation in floral morphology, pollinator morphology and behaviour, location of pollen on the pollinator’s body, and other temporal and spatial factors besides the quantity of pollen transported. Instead, interaction frequency (counting occurrences of an interaction, but disregarding individual interaction strength) predicts the relative strength of pollination interactions well (Vázquez *et al.*, 2005), and was successfully generated with both microscopy and metabarcoding in our study.

### Merging metabarcoding and pollination network analysis

Following several recent studies which have constructed diurnal plant-pollinator networks using DNA metabarcoding (Bell *et al.*, 2017; Pornon *et al.*, 2017), we have further demonstrated the potential of metabarcoding by using it to construct nocturnal pollen-transport networks for the first time (Fig. 3). We provide a detailed and reproducible methodology to integrate molecular advances and ecological network analysis. Our results clearly demonstrate that the capacity of metabarcoding to generate pollen-transport interaction data is comparable to that of previously-used methods, such as microscopy. Additionally, metabarcoding may carry several practical advantages over flower-visitor observations or microscopy for studies analysing pollination networks.

One such advantage is that metabarcoding is reproducible across studies, pollinator guilds, and ecosystems. It is freed from observer biases inherent both in morphological identification of pollen, and in other means of detecting pollination interactions such as flower-visitor observations, where distribution of sampling effort among flower species can affect network structure (Gibson *et al.*, 2011) and sampling often focuses on a subset of the floral assemblage (e.g. Tiusanen *et al.*, 2016). Metabarcoding can be conducted without system-specific expertise in morphological pollen identification, or prior knowledge about locally-present plants or likely interactions (although such information can be used, if available and robust, to increase the taxonomic resolution of species identifications). Metabarcoding may reveal previously unsuspected detail in networks (Pornon *et al.*, 2017), especially those involving moths or other under-studied pollinator taxa.

Metabarcoding may also allow more efficient processing of samples, and therefore the analysis of larger numbers of samples, than microscopy (Fig. 5). Most pollination-network studies have focused on evaluating a single network, or a small number of networks under variant conditions (e.g. Burkle *et al.*, 2013). Constructing multiple replicated networks across a range of treatments, sites or time points, and testing for structural differences (e.g. Lopezaraiza-Mikel *et al.*, 2007), is a powerful alternative, but can be hampered by the difficulty of generating enough data for multiple, well-sampled networks. For metabarcoding, investment mainly scales per-plate (≤ 96 samples) rather than per-sample (Derocles *et al.*, 2018), whereas for microscopy, investment of materials and especially time increases linearly for every sample, although sample-processing speed might increase slightly after an initial period of learning (Fig. 5). Importantly, this increased efficiency is coupled with increased reproducibility, as molecular tools treat all samples identically regardless of their complexity.

**Figure 5:**
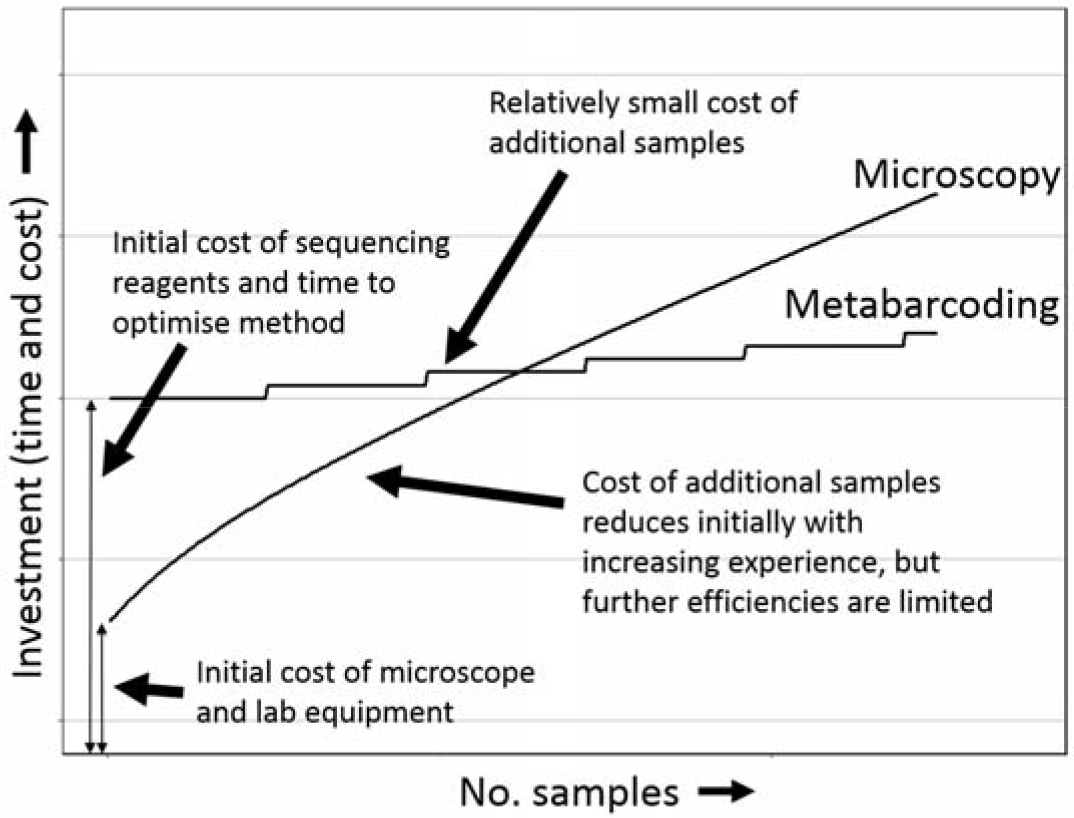
**estimated change in investment as number of samples increases** for metabarcoding and microscopy methods. Lines are hypothetical and not based on formal costing of methods.

Finally, DNA metabarcoding can streamline the generation of suitable data for incorporating phylogenetic information into ecological networks (Evans *et al.*, 2016). Recent studies have found significant relationships between phylogenetic and resource overlap in mutualistic and antagonistic networks (Rezende *et al.*, 2007; Elias *et al.*, 2013; Peralta *et al.*, 2015); metabarcoding permits simultaneous generation of both interaction and relatedness data.

### Conclusions

In this study, we constructed pollen-transport networks using matched samples of moths to compare between two methods for detecting and identifying pollen: DNA metabarcoding and traditional light microscopy. We showed that the state-of-the-art DNA metabarcoding approach is capable of generating pollen-transport interaction networks that are similar to those detected using microscopy. Indeed, with metabarcoding, we detected pollen on more individual moths and detected more pollen types per moth species. These differences indicate that direct comparisons between networks constructed using metabarcoding and those constructed using traditional methods such as microscopy should be treated with appropriate caution, but a combination of both metabarcoding and traditional methods may provide the most detailed information (Wirta *et al.*, 2014). Metabarcoding additionally revealed a range of previously undocumented moth-plant interactions, and provided new evidence for two possible benefits of nocturnal pollination: landscape-scale provision of plant gene flow, and potential provision of the pollination ecosystem service. The metabarcoding approach has considerable potential for studying pollen-transport networks and species-interactions more generally.

## Acknowledgements

This work was supported by the Natural Environment Research Council and Butterfly Conservation (Industrial CASE studentship awarded to C.J.M., Project Reference: NE/K007394/1) and was conducted with ethical approval from the University of Hull (Approval Code U074). We thank T. Hall for her permission to sample moths at Molescroft Grange Farm. We thank A. Lucas and N. de Vere for useful discussions prior to commencing labwork, and J. Downs for assistance with fieldwork. E. Moss created the moth image in Fig. 1.

## Contribution of authors

The experiment was conceived by C.J.M. under supervision by D.M.E., M.J.O.P and R.F. and designed by those authors with D.H.L. and J.J.N.K. Field and laboratory work was conducted by C.J.M. with advice from J.J.N.K. The metaBEAT pipeline was created by C.H. and metabarcoding data was processed and analysed by C.J.M., with advice from C.H. The statistical analysis was conducted by C.J.M. All authors contributed to preparing the manuscript and gave final approval for publication.

## Data Accessibility Statement

- Raw DNA sequence reads: Sequence Read Archive, accession number SRP102977.
- Bioinformatic and analytical scripts: Zenodo, doi: 10.5281/zenodo.1169319.
- Processed interaction data: Dryad doi: …(upon acceptance)

